# OptiCell3D: Precise inference of mechanical cell properties from microscopy imaging

**DOI:** 10.64898/2026.05.23.727231

**Authors:** Steve Runser, Kevin A Yamauchi, Marius Almanstötter, Franziska L Lampart, Federico Carrara, Lisa Conrad, Laura Schaumann, Roman Vetter, Dagmar Iber

**Author notes:** These authors contributed equally to this work (Steve Runser, Kevin A Yamauchi).

## Abstract

We introduce OptiCell3D, an open-source image-based framework for inferring cellular mechanical properties directly from 3D microscopy data. By integrating short physical simulations with gradient-based optimization via backpropagation, our method substantially improves accuracy compared to existing approaches. Moreover, OptiCell3D enables mechanical inference in tissues that are too large to be fully imaged, extending its applicability to more complex biological systems. We demonstrate the power of our approach by applying OptiCell3D to five morphologically diverse mouse epithelial tissues. By combining inferred parameters with simulations and morphometric analysis, we find that the ratio of apical to lateral surface tension predicts cell aspect ratio across epithelial subtypes, linking a single mechanical parameter to the broad morphological diversity of epithelia. Finally, we apply our framework to stratified tissues, finding greater variability in pressure and surface tension between cells and tension gradients along the apico-basal axis.

## Introduction

Tissue morphogenesis and disease progression are governed by mechanical forces that act on the scale of individual cells. Quantifying these forces and the underlying mechanical properties is therefore central to understanding how tissues are built and how they fail. Established experimental methods such as atomic force microscopy, deformability cytometry, and micropipette aspiration measure the mechanical properties of cells and tissues by probing cell deformation under a controlled load, but require direct physical access to the sample and are therefore poorly suited for measurements deep within intact tissues or living organisms [1]. Particle-Tracking Microrheology, Magnetic tweezers, and embedded oil droplets extend mechanical measurements into tissue interiors by embedding exogenous materials, but this can perturb the mechanical environment they aim to measure [2–4]. Brillouin microscopy can map mechanical properties optically without exogenous labels or embedded materials, but its relationship to rheologically defined moduli remains an active area of investigation [5–7].

Inferring forces directly from cell morphology has emerged as a promising alternative, requiring no physical contact or exogenous materials. Early image-based inference methods were limited to analyzing cell surface tensions and pressures in 2D tissue slices [8–16]. Recent advances now permit force inference from 3D tissue geometries [17–23] (Table 1). Regardless of dimensionality, all image-based methods rely on one of two approaches for estimating cell pressures and surface tensions. The first one mechanically links measured tri-cellular junction angles and cell interface curvatures. Cell contact angles are traditionally measured manually or automatically in images [24, 25], but limited image resolution and the fact that cell boundaries span multiple pixels can lead to significant errors in angle estimation. The second method, known as the variational approach, discretizes the tissue geometry using a mesh, and then optimizes parameters to balance forces at its nodes, enabling estimation of additional parameters beyond surface tensions and pressures. The variational approach can be further refined by finding parameters that keep mesh nodes in place during a short mechanical simulation rather than balancing forces only in the input geometry [19]. This subtle difference enhances the method’s resilience to inaccuracies in the input data. Here, we introduce a new framework called OptiCell3D that builds on this idea by integrating it with differentiable numerical simulations [15], thereby further enhancing inference accuracy and performance. Moreover, we incorporate the ability to treat selected segmented tissue boundaries as static during parameter inference, which improves the accuracy of inferred parameters for tissues that are too large to be fully imaged. To demonstrate its applicability, we apply OptiCell3D to different epithelial tissues.

Epithelia are continuous sheets of tightly connected cells lining body surfaces that display a remarkable diversity of cell morphologies across tissues and organisms [26]. They are organized either as a single layer (simple epithelium) or multiple layers (stratified epithelium), with individual cells ranging from squamous and cuboidal to columnar and pseudostratified forms. Biophysical models have linked the cell aspect ratio to the apical-to-lateral surface tension and cell-substrate adhesion [27]. How strongly the underlying biophysical parameters vary to support these different geometries remains largely unknown. In budding intestinal organoids, apical-to-lateral and basal-to-lateral tension ratios of less than 20% have been inferred, suggesting that even minor tension differences can induce relevant morphological tissue changes [23]. Therefore, high-precision force inference methods are required to understand how the rich morphospace of epithelia emerges. By applying OptiCell3D across multiple epithelial tissue types, we provide the first systematic quantitative comparison of how two key biophysical parameters, surface tension ratios and cell pressures, vary across this morphological diversity. We quantify how the cell aspect ratio increases with the ratio of apical to lateral surface tension and confirm this relationship in SimuCell3D simulations [28], linking a simple mechanical parameter to the broad morphological diversity observed across epithelia. Extending this analysis to stratified tissues, we find that both surface tensions and cell pressures show greater cell-to-cell variability than in simple epithelia, and observe a decrease in mean cell surface tension but not cell pressure along the apical-basal axis.

## Results

### OptiCell3D

OptiCell3D infers cellular forces from 3D images by using gradient-based optimization to iteratively determine the mechanical properties that best reproduce the observed cell geometries, assuming mechanical equilibrium (Fig. 1a). A differentiable deformable cell model is parameterized by the intracellular pressure of each cell and the surface tension of each interface (Fig. 1b, Methods). Implemented in JAX [29], OptiCell3D is automatically differentiable and can be accelerated on a variety of hardware (e.g., graphics and tensor processing units). The parameter inference pipeline takes a cell surface mesh and a user-specified parameter range as input. In each iteration of the optimization, a short mechanical simulation is run with overdamped dynamics using a random initial parameter guess (Methods). Next, a loss function based on the obtained nodal displacements (Eq. 11) is iteratively minimized to find better parameter estimates. The optimization is terminated after a user-specified number of iterations, and repeated with a new random initial guess for a user-specified number of times. This multi-start optimization strategy reduces sensitivity to local minima. The parameter set with the lowest loss is taken as the best estimate. The returned cell pressures and interfacial tensions are non-dimensionalized and normalized so that their arithmetic mean value vanishes.

**Figure 1:**
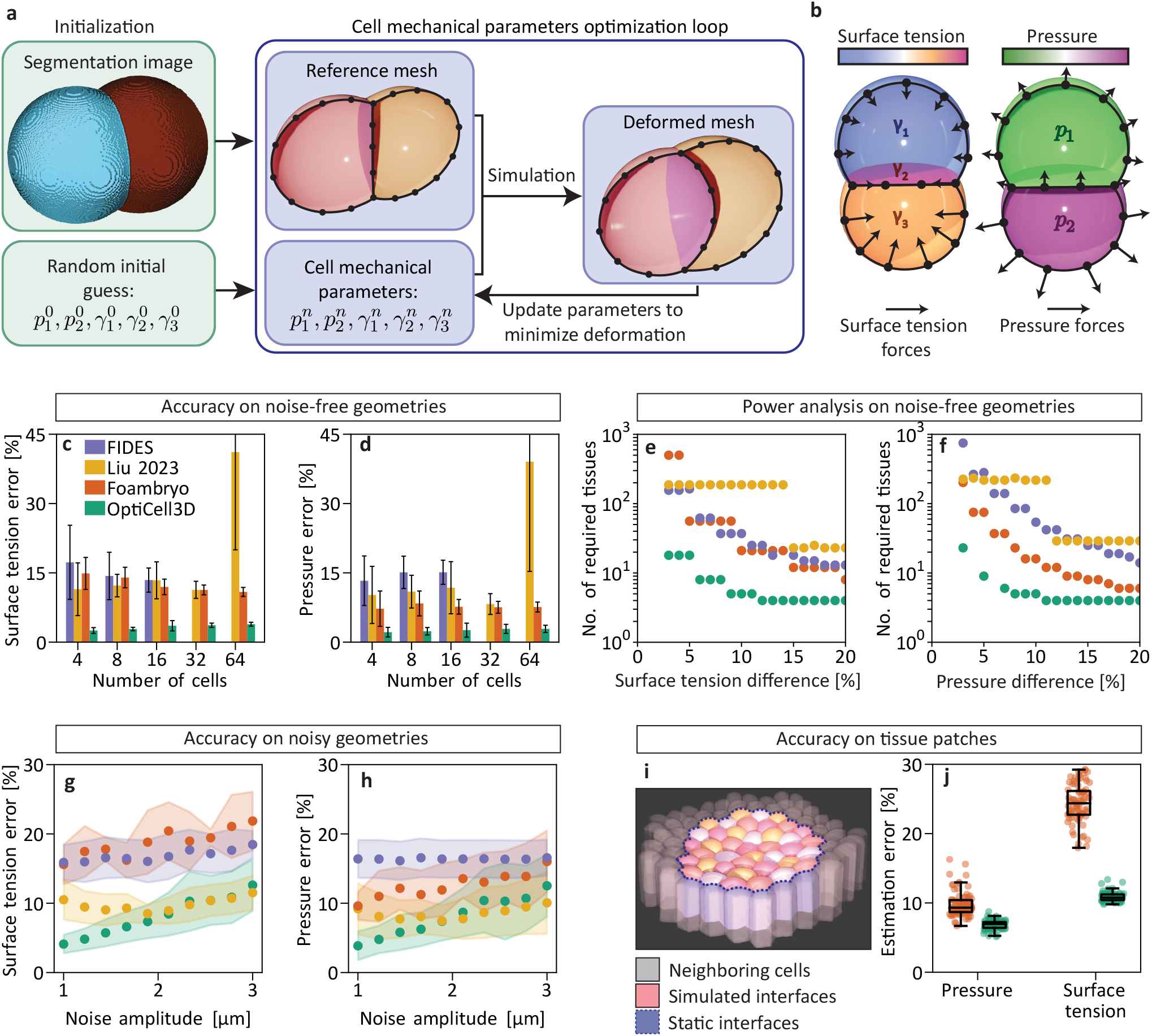
Comparison of cell parameter inference methods. **a** Illustration of the gradient-based optimization pipeline of OptiCell3D to infer individual net pressures *p* and surface tensions *γ* from 3D images. **b** In mechanical equilibrium, forces from cell-internal hydrostatic net pressure and interfacial surface tension are balanced. **c**,**d** Benchmark of inference accuracy of OptiCell3D, Foambryo [18], FIDES [19], and “Liu 2023” [20] using *n* = 50 simulated geometries. 32 and 64-cell tissues were excluded for FIDES due to prohibitively long runtimes (> 15 days). **e**,**f** Number of samples required to detect a relative parameter difference at a 95% confidence level. **g**,**h** Inference accuracy on *n* = 10 8-cell geometries perturbed with varying amplitudes of Perlin noise using 1 octave. **i** Example geometry used to benchmark inference performance on patches of tissue, with fixed tissue boundaries (cell-cell interfaces) in blue. **j** Inference accuracy on *n* = 80 patches of tissue. All error bars represent SD over the *n* geometries. Colors in panel c apply to all data panels.

To investigate how the inference error of OptiCell3D scales with the number of cells in a tissue, we generated synthetic geometries (Methods) with known cell pressures and surface tensions containing 4, 8, 16, 32, and 64 cells (*n* = 250 geometries in total). We then quantified the surface tension and pressure inference errors of OptiCell3D and compared them to those obtained using three other 3D image-based inference methods: Foambryo [18], FIDES [19], and the method developed by Liu *et al*. (hereafter referred to as “Liu 2023”) [20] (Fig. 1c,d). The estimation error showed no significant dependence on cell number for any method except “Liu 2023”, for which the error increased markedly from tissues with fewer than 64 cells to those with 64 cells, rising from 10.29 ± 4.83% (mean ± SD) to 39.05 ± 23.73% for pressure and from 12.10 ± 3.86% to 41.13 ± 21.15% for surface tension. OptiCell3D achieved estimation errors of 2.32 ± 0.72% for pressure and 3.11 ± 0.58% for surface tension across all tissue sizes. In comparison, the mean estimation errors of FIDES and Foambryo were 3.5-to 7.2-fold higher. OptiCell3D also had consistently narrower error distributions than the other tested methods (Supplementary Fig. S1).

Sample size determination is crucial for any empirical study aimed at drawing conclusions about observed populations. We evaluated how OptiCell3D reduces the sample size needed to detect parameter differences with statistical confidence. We generated synthetic parameter populations by bootstrapping the error distributions (Supplementary Fig. S1) obtained from the unperturbed geometry benchmark (Fig. 1c,d) and statistically compared them. For each method tested, the *p*-value for a desired parameter difference decayed with the number of tissues at different rates (Supplementary Fig. S2). OptiCell3D requires over 4 times fewer tissues than Foambryo to detect a 10% difference in surface tension at a 95% confidence level, 7 times fewer than FIDES, and 37 times fewer than “Liu 2023” (Fig. 1e). For the same difference in pressure, we observed a reduction in the required sample size by over 3 times compared to Foambryo, 11 times compared to FIDES, and 43 times compared to “Liu 2023” (Fig. 1f).

Due to noise or artifacts in microscopy images and errors introduced by segmentation methods, the positions of cell boundaries provided to image-based inference tools are often inaccurate. To evaluate the impact of this error on the parameter inference, we produced a set of *n* = 10 8-cell geometries perturbed by Perlin noise [30] (Methods and Supplementary Fig. S3) with controlled noise amplitude and frequency. A low noise frequency was used to generate realistic perturbations mimicking imaging inaccuracies. OptiCell3D largely outperformed Foambryo and FIDES across all tested noise intensities (Fig. 1g,h). At high noise intensity, OptiCell3D and “Liu 2023” exhibited the same error levels. A full analysis with varying noise intensities and frequencies shows the expected increase in accuracy at low noise levels for OptiCell3D, but rather uniform errors for Foambryo, FIDES, and “Liu 2023” (Supplementary Fig. S4).

Current 3D parameter inference algorithms are designed for tissues that are fully visible within the segmented image. This limitation prevents inference in situations where imaging the entire tissue is not possible (e.g., epithelial sheets). We have removed this limitation by introducing boundary conditions that allow the user to make specified interfaces static. To demonstrate the applicability of OptiCell3D to incomplete tissue images, we generated *n* = 80 tissue sheets composed of adherent columnar cells using SimuCell3D [28] and selected subsets of their cells for parameter inference (Fig. 1i). OptiCell3D achieved pressure and surface tension errors on these tissue patches of 6.95 ± 0.78% and 10.85 ± 0.62%, respectively (Fig. 1j). Foambryo recorded errors of 9.81 ± 1.79% for pressure and 24.38 ± 2.75% for surface tension, which are 1.42 and 2.25 times as large.

We also measured the real (wallclock) time required to convert an input image into a mesh and estimate the cell parameters from it. For 8-cell tissues, FIDES required 6.5 hours to perform this task, “Liu 2023” required 28 seconds, while OptiCell3D completed it in 2 minutes, and Foambryo took only 4 seconds on an ordinary workstation (Supplementary Fig. S5).

### 3D morphometric analysis of cells in diverse epithelial tissues

To gain insight into the mechanical properties underlying diverse epithelial architectures and to demonstrate the scope and accuracy of OptiCell3D, we applied the framework to high-resolution 3D light-sheet fluorescence microscopy images of five distinct epithelial tissues from mouse embryos: bronchiole, kidney, intestinal villus, esophagus, and bladder. These tissues span a wide range of cellular geometries, from cuboidal to columnar, and structural organizations, from single-layered to stratified epithelia (Fig. 2a).

**Figure 2:**
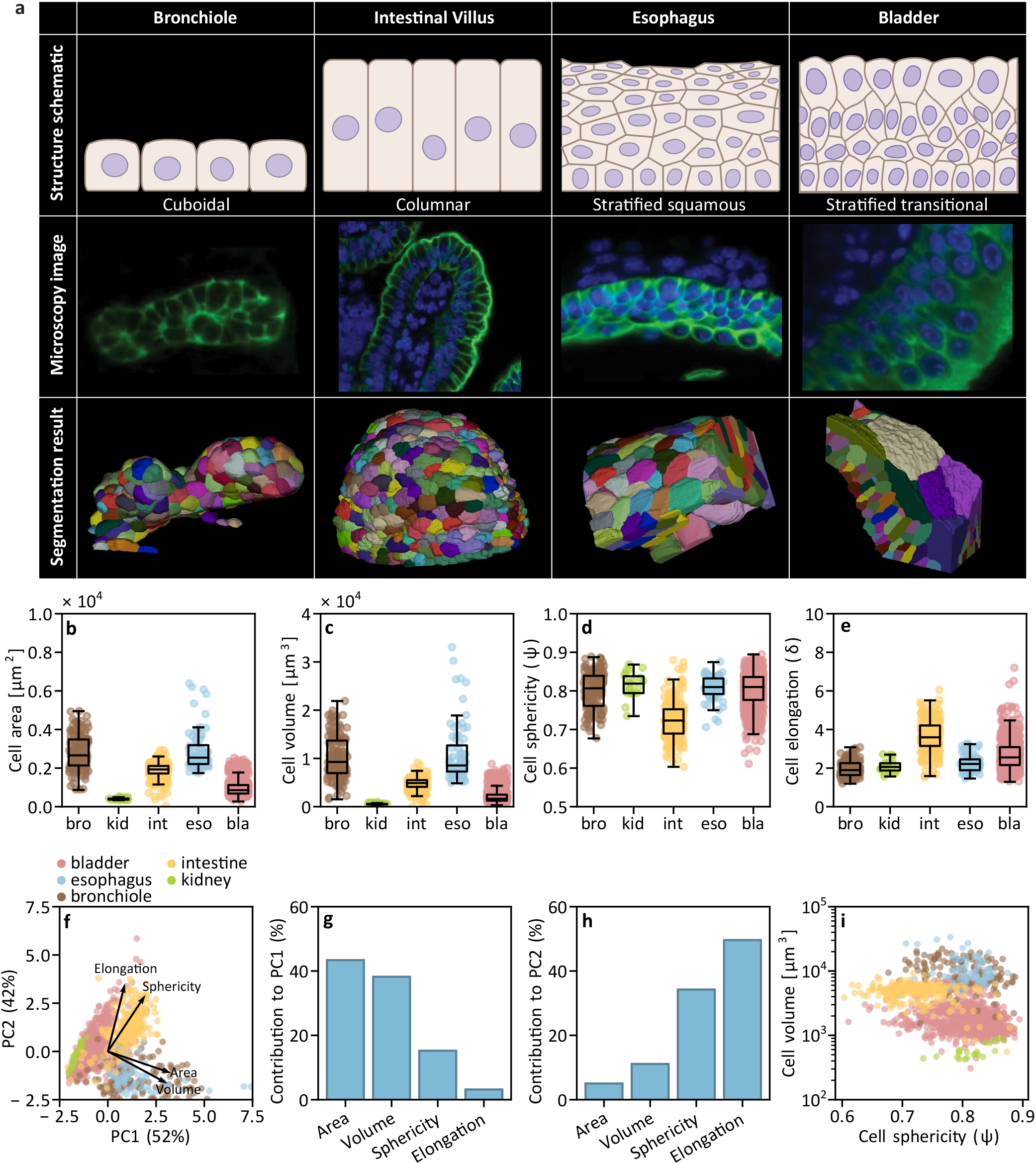
Morphometric analysis of cells across epithelia. **a** Top row: 2D schematic illustrating epithelial cell shape and tissue organization in the five epithelial architectures analyzed in this study: cuboidal epithelium in the bronchiole and kidney, columnar epithelium in the intestinal villus, stratified squamous epithelium in the esophagus, and transitional epithelium in the urinary bladder. Middle row: Light-sheet microscopy images of the different epithelia, showing cell membranes in green and nuclei in blue. Bottom row: Corresponding 3D segmentation, with individual cells color-coded by label. **b–e** Distribution of geometric features for bronchiole, kidney, intestinal villus, esophagus, and bladder tissues. Boxplots represent the interquartile range (IQR), with the central line indicating the median and whiskers extending to the most extreme data points within 1.5 times the IQR. Individual data points are overlaid to show the underlying sample distribution and density. **f** Principal component analysis based on the extracted cell morphological features. The first two principal components (PC1, PC2) account for 52 and 42% of the total variance, respectively. Overlaid vectors represent the original features; the direction and length of each vector indicate its contribution (loading) to the principal components. **g**,**h** Contribution in percentages of each morphological feature to the first and second principal components. **i** Scatter plot displaying the distribution of cell volumes relative to sphericity across the five tissue types.

We started by analyzing the morphologies of the segmented cells to quantify variations in cell size and shape across the five tissue types. This morphometric analysis revealed that the cells in the kidney were markedly smaller than all other cell types (Fig. 2b,c and Supplementary Fig. S6a,b). The bronchiole and esophagus exhibited statistically comparable volumes and surface areas, both significantly larger than those of cells in the intestinal villus, kidney, and bladder. Intestinal villus cells were significantly less spherical than those in the other tissues (Fig. 2d and Supplementary Fig. S6c), consistent with their characteristic columnar morphology. Cells from the intestinal villus and bladder displayed significantly greater elongation than those from the bronchiole, kidney, and esophagus (Fig. 2e and Supplementary Fig. S6d). The sample size for each tissue type, along with the mean and standard deviation of the features analyzed here, are summarized in Table 3.

Principal component analysis indicates that variation across these tissues arises primarily from differences in cell size, followed by shape anisotropy (Fig. 2f–h). Overall, kidney cells were substantially smaller than those of all other tissues. Cells from the bladder and intestinal villi were also smaller and more anisotropic than those from the bronchioles and esophagus (Fig. 2i).

### Cell height control in monolayer epithelial tissue

To investigate the mechanical origins of the differences in cell shape observed in monolayer epithelia, we applied our framework to bronchiole, kidney, and intestinal tissues (Fig. 3a). In all monolayer tissues, we observed significantly higher surface tensions at the apical and basal surfaces compared to the lateral surfaces (Fig. 3b). Given this major difference, we hypothesized that the ratio of apical to lateral surface tension (*α*) plays a key role in determining cell shapes in monolayer tissues. To test this hypothesis, we computed the surface tension ratio and the aspect ratio for each segmented cell (Fig. 3c,d). In parallel, we performed mechanical forward simulations using SimuCell3D [28], in which cells were allowed to deform and slide over a fixed substrate until reaching mechanical equilibrium (Fig. 3c). In these simulations, we prescribed different values of *α* and measured the resulting cell aspect ratios at steady state. Both experimental and simulation results show that increasing *α* leads to greater cell aspect ratios (Fig. 3e). The deviation at high alpha values between experimental and simulated results may arise from boundary effects associated with the finite size of the simulated patch. Taken together, these findings indicate that relative changes between apical and lateral surface tensions can drive the transition from cuboidal to columnar cell shapes in simple epithelia.

**Figure 3:**
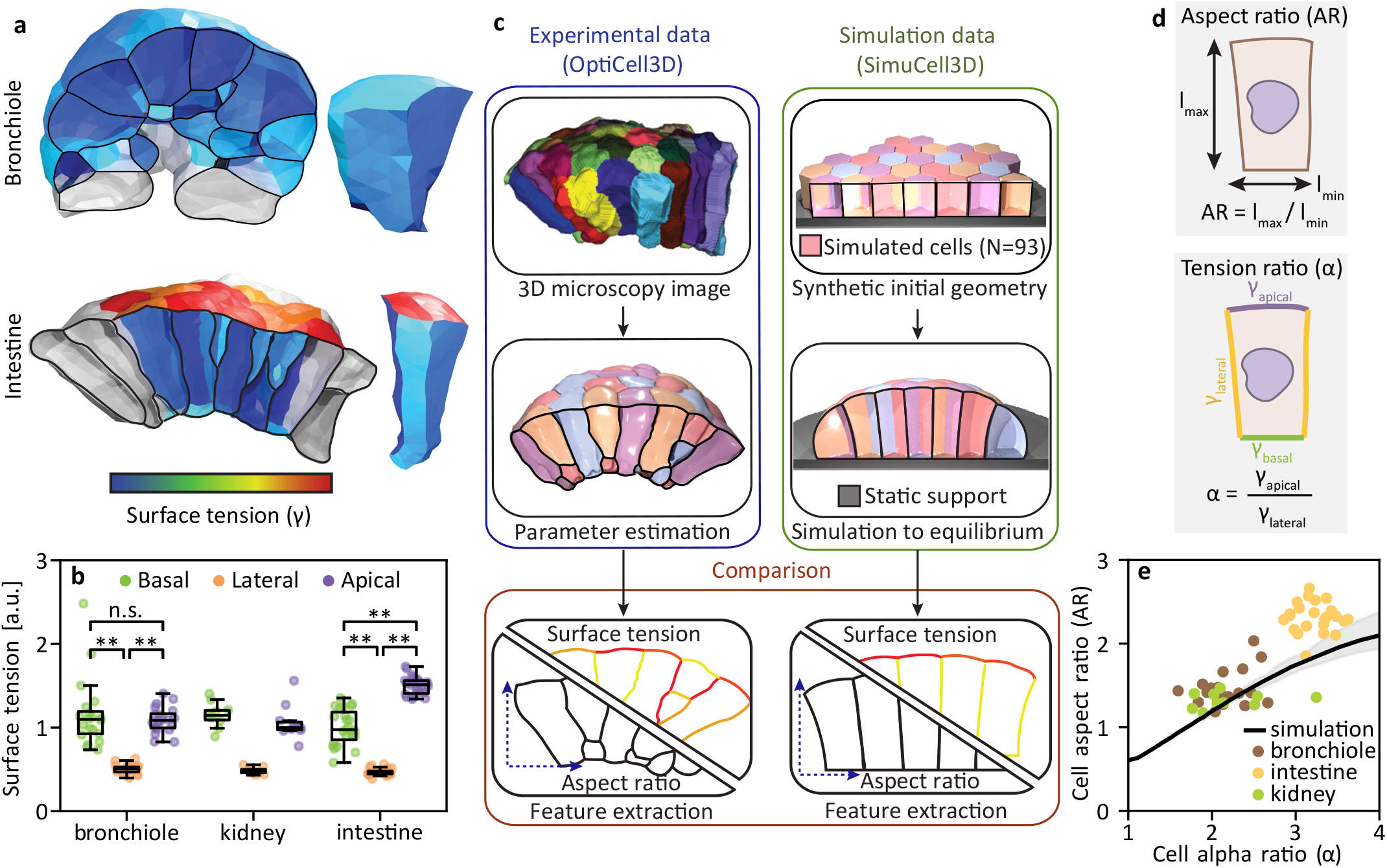
Surface tension distribution in monolayer epithelial tissues. **a** Left: 3D renderings of analyzed bronchiole and intestine tissue patches. Right: Representative 3D renderings of individual cells from each tissue, with interfaces color-coded by inferred surface tension. **b** Distribution of surface tensions across basal, lateral, and apical interfaces in bronchiole, kindey, and intestine epithelia. Stars indicate the significance of differences between distribution means. **c** Analysis pipeline relating the apico-lateral surface tension ratio *α* to cell aspect ratio (AR) in experimental data and forward simulations. Experimental patches are extracted from 3D segmentations, meshed, and processed using OptiCell3D to estimate surface tensions. Simulated tissues with prescribed *α* values are generated with SimuCell3D [28], with the AR measured in mechanical equilibrium. **d** Top: Calculation of the AR via an axis-aligned bounding box. Bottom: Definition of the apico-lateral surface tension ratio *α* = *γ*_apical_*/γ*_lateral_. **e** Comparison of experimental and simulated trends, indicating that increased *α* correlates with increased cell height in monolayer epithelia. Shaded gray regions indicate the minimum–maximum range of observed aspect ratios within each simulated patch. Boxplots indicate the median (center line) and interquartile range (IQR), with whiskers extending to the most extreme data points within 1.5 × IQR. Significance level: \*\**p* < 0.01. *p*-values were computed using a permutation test on CLR-transformed data (Methods) and then adjusted for multiple comparisons using the Benjamini–Hochberg procedure. Pairwise tests were performed only when both groups contained at least 15 samples.

### Apico-basal tension gradients in stratified tissues

Next, we applied OptiCell3D to stratified epithelia, which revealed that variability in mean cell surface tension and pressure is significantly greater in stratified tissues than in monolayer epithelia (Fig. 4a,b). To assess whether this variability correlates with cellular position, we assigned a layer index to each cell using a depth-first search on the cell–cell adjacency graph, starting from the basal surface (Fig. 4c). Plotting mean cell surface tension as a function of layer index revealed a clear increase between the first and second layers in both stratified tissues (Fig. 4d). In contrast, pressure variability did not appear to arise from systematic differences across layers, as no significant variation in pressure was observed between them (Fig. 4e). To further dissect the observed trend in surface tension, we asked whether the increase along the apico–basal axis reflects a global shift across all interface types. In the esophagus, we observed a significant decrease in surface tension between basal and lateral interfaces within the first layer, followed by an increase across subsequent layers (Fig. 4f). However, the limited number of inferred measurements precluded robust statistical testing of this trend. In the bladder, surface tension increased significantly from the basal interface of the first layer to the lateral interface of the second layer, after which it reached a plateau with no further significant increases in upper layers (Fig. 4g,h). Overall, these results indicate that in stratified tissues, surface tension increases from the basal side to the apical side within the first layer and then remains approximately constant across subsequent layers.

**Figure 4:**
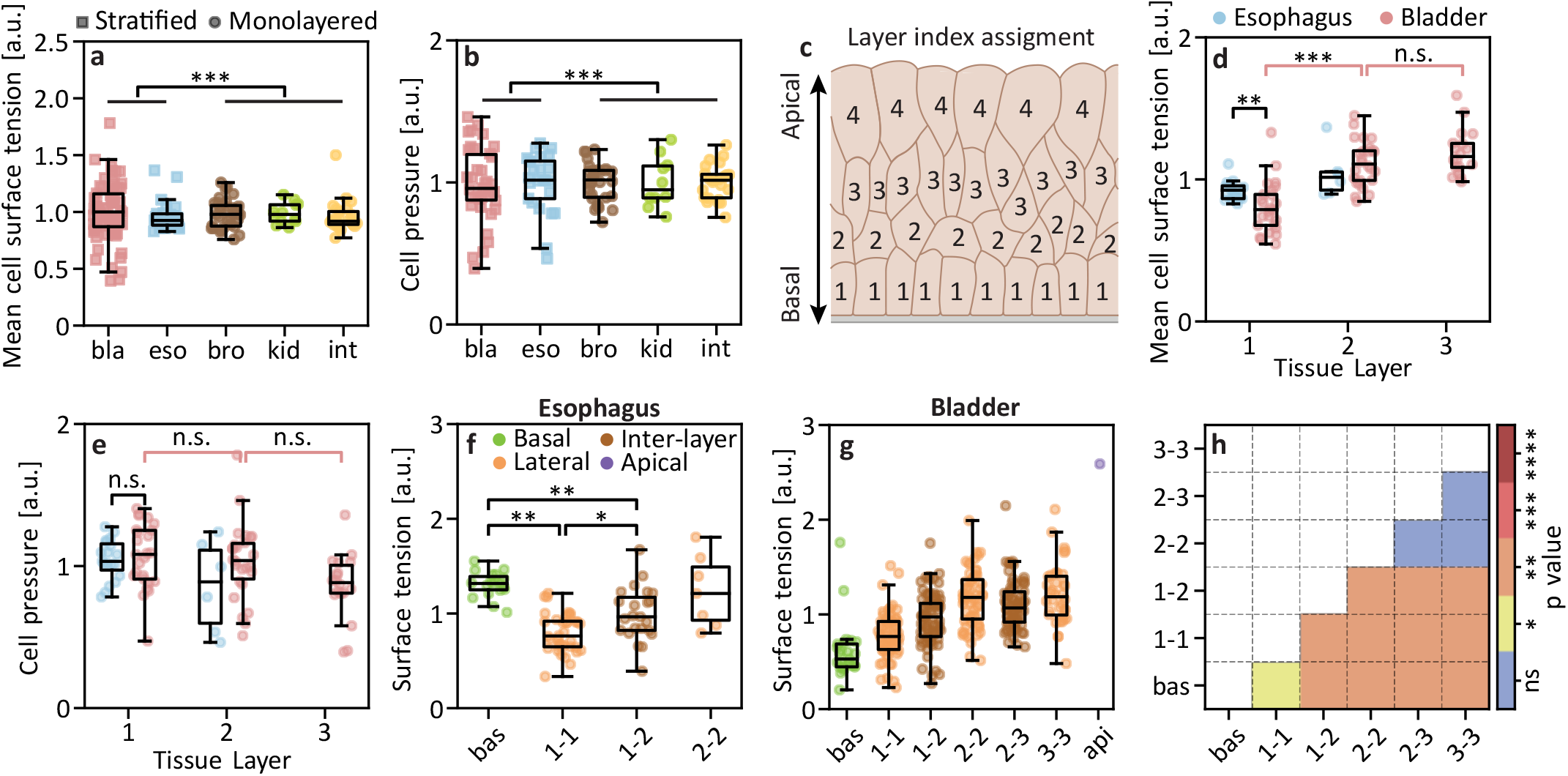
Surface tension distribution in the analyzed stratified epithelial tissues. **a**,**b** Boxplots showing the distribution of mean cell surface tension (**a**) and pressure (**b**) across the five analyzed epithelial tissue types, with individual data points overlaid. Significant difference in variance between stratified (bladder, esophagus) and monolayered (bronchiole, kidney, intestinal villus) tissues are indicated by three asterisks. **c** Schematic illustrating the assignment of layer indices to individual cells based on their topological distance from the basal surface. **d**,**e** Mean cell surface tension (**d**) and pressure (**e**) as a function of layer index in stratified tissues. **f**,**g** Distribution of surface tension across basal, lateral, inter-layer, and apical interfaces in esophageal and bladder epithelia. **h** Pairwise comparison of surface tension across bladder interfaces. Boxplots show the median (center line) and interquartile range (IQR; box limits), with whiskers extending to the most extreme data points within 1.5 × IQR. Significance levels: \**p* < 0.05, \*\**p* < 0.01, and \*\*\**p* < 0.001. *p*-values were computed using a permutation test on CLR-transformed data (Methods) and adjusted for multiple comparisons using the Holm–Bonferroni procedure. Unless otherwise stated, the mean was used as the test statistic for comparing distributions. Pairwise tests were performed only when both groups contained at least 15 samples.

## Discussion

We have introduced OptiCell3D, a computational open-source framework for mechanical parameter inference from 3D microscopy images that achieves improved accuracy over existing approaches across a wide range of tissue sizes and imaging noise levels. This increase in accuracy substantially reduces the number of samples required to detect differences in cellular properties between experimental conditions. Since cell morphology can be sensitive to small changes in interfacial tension differences, such high-precision quantifications are required to understand the wealth of epithelial tissue architectures observed in nature, and how they emerge and transition into each other. Current 3D inference methods have been primarily applied to early mammalian embryos, where inference has revealed only stochastic variability rather than systematic differences between cell fates so far [21], leaving the question whether molecular or mechanical cues drive cell internalization open [31]. The improved accuracy of OptiCell3D may help resolve such questions by reducing inference noise below biologically meaningful signal levels. We further demonstrated the applicability of our method on tissue patches that are too large to be fully segmented, thereby extending the applications of image-based inference methods to a broader set of tissues.

By applying OptiCell3D to segmented images of epithelial tissues, we have gained several insights into the mechanical determinants of their architecture. First, consistent with theoretical predictions [27], our results suggest that the ratio of apical to lateral surface tension controls the shape of cells within epithelial monolayers. The quantitative range of *α* values we observe across tissues implies that modest changes in actomyosin contractility or cell-cell adhesion, the two main molecular determinants of interfacial tension, may be sufficient to drive large-scale morphological transitions between epithelial subtypes. However, the cuboidal-to-squamous transition in the *Drosophila* wing disc has been linked to mechanical confinement arising from tissue growth [32]. Static image-based inference alone cannot distinguish between these mechanisms. Rather, time-lapse inference and perturbation experiments would be required. Second, applying OptiCell3D to stratified epithelia revealed higher variability in the mean cell surface tension and pressure as well as a gradient of surface tensions along the transverse axis of the tissue, concentrated between the basal and adjacent layers, with tensions remaining approximately constant in more apical layers. Whether this variability is a cause or consequence of stratification remains an open question that future studies combining mechanical inference with live imaging and perturbation experiments could address.

These results were obtained using a deliberately simple mechanical description of cells, limited to intracellular net pressure and interfacial surface tensions. This formulation could be readily extended to incorporate additional mechanical behavior, such as hyperelasticity, to better capture the complexity of real tissues. A central limitation of the current approach, however, is that the mesh topology is fixed during the short differentiable simulations, preventing cell rearrangements and the formation of new contacts. Relaxing this constraint would significantly broaden the scope of the method. Beyond inferring mechanical parameters from static image snapshots, as performed here, it would enable prediction of how these parameters must evolve over time to drive tissue transitions between distinct topologies. Such capabilities may, for example, pave the way to applications of control theory in tissue development, where control policies are first optimized on a virtual tissue twin before being applied experimentally to guide morphogenesis. Mesh rearrangements were not implemented in OptiCell3D because they introduce discontinuities in gradient computations, which compromise differentiability. However, recent advances in simpler two-dimensional differentiable cell-based models have demonstrated robust strategies to handle these discontinuities while allowing for topological transitions [15]. Extending such approaches to more geometrically detailed models, such as OptiCell3D, represents a promising direction toward fully differentiable simulations of multicellular systems. An alternative strategy would be to shift from a Lagrangian (mesh-based) representation of tissue geometry to an Eulerian (grid-based) formulation, such as the one used in phase-field or level-set methods [33–35]. In such formulations, cell rearrangements emerge naturally without requiring explicit topological updates, thereby avoiding discontinuities altogether.

Overall, we anticipate that differentiable simulations of high-resolution, cell-based models will become increasingly prevalent in the coming years. We believe these advances have the potential to enable a new paradigm for investigating the relationship between cellular mechanical properties and the emergent tissue morphologies.

## Methods

### Ethics statement

Use of animals was approved by Canton Basel-Stadt (license number 2777/33495). Experimental procedures were performed in accordance with the Guide for the Care and Use of Laboratory Animals and approved by the Ethics Committee for Animal Care of ETH Zurich. All animals were housed at the D-BSSE/UniBasel facility under standard water, feeding, enrichment, and 12-hour light/dark cycles.

### Mouse model

Ureteric buds were isolated from E12.75 transgenic Mouse: Tg(Hoxb7-Venus^∗^)17Cos embryos that express a membrane-associated form of the fluorescent protein Venus under the control of the *Hoxb7* promoter [36]. Adult mouse bladder data were reused from Lampart et al. [37]. All other epithelial tissues were dissected from E17.5 mouse embryos homozygous for Rosa26mTmG and heterozygous for the ShhGFP-Cre allele (ShhGFP-Cre/+; Rosa26mTmG). The epithelial cell membranes are labeled with GFP following CRE-mediated excision [38]. Offspring from Shhcre x mT/mG breedings show membrane-targeted tdTomato (mT) in all tissues, but have the mT cassette deleted for tissues expressing *Shh* and therefore express membrane-targeted EGFP (mG).

### Immunofluorescence staining and optical clearing

The isolated tissues were fixed in 4% paraformaldehyde (PFA) in PBS (28908, Thermo Fisher, brand name) for 1.5 hours at 4 deg. C. Samples were washed 4 times with PBSTx (0.25% Triton X-100 in PBS) following fixation. Samples were incubated with CUBIC-1 clearing solution for 3 days at 37 deg C. Following the first clearing step, samples were blocked for 2 days in 1% BSA (catalog number, brand name A7906, Sigma), 10% goat serum (S26-100ml, Sigma) in PBSTx at 4 deg. C on a shaker. The samples were incubated with the primary antibodies, anti-GFP (1:500, GFP-1020, aves) and anti-laminB1 (1:200, 702972, Thermo Fisher), for 2 days at 4 deg. C. The primary antibodies were diluted in blocking buffer. After primary antibody incubation, the samples were washed 3 times with PBSTx. The samples were incubated with secondary antibodies, goat anti-chicken conjugated with Alexa Fluor 488 (1:200,A11039, Invitrogen), donkey anti-rabbit conjugated with Alexa Fluor 647 (1:200, A32795, Invitrogen), and DAPI (1:2000, 1mg/ml stock solution, D9542, Sigma) for 3 days at 4 deg. C. The secondary antibodies were diluted in blocking buffer. After secondary antibody incubation, the samples were washed 3 times for 15 minutes in PBSTx at 4 deg. C. Samples were fixed a second time in 4% PFA in PBS for 30 minutes at 4 degrees C. Samples were washed 4 times in PBSTx for 15 minutes at 4 deg. C. The samples were incubated in 50% CUBIC-2 solution diluted in ddH2O for 1 day. Then they were then incubated in 100% CUBIC-2 until fully cleared (depending on sample size). During the 100% CUBIC-2 incubation, the CUBIC-2 solution was exchanged every other day. Finally, they were immersed in a freshly-prepared CUBIC-2 solution. All CUBIC-2 incubations were performed at room temperature.

### Imaging of whole-mount epithelial tissues

Cleared and stained samples were embedded in 2% agarose cylinders and incubated in CUBIC-2 for 2 days. Images were acquired with a Zeiss Z.1 lightsheet microscope using a 20X 1.0NA Plan Neofluar objective (421459-9970, Zeiss).

### 3D morphometry of cells in epithelial tissues

#### 3D cell segmentation

To extract the geometries of the individual cells in the tissues, we performed 3D instance segmentation using Cellpose. We performed the 3D segmentation by predicting flows on 2d slices in the X, Y, and Z directions via the do 3d option. As inference of mechanical properties relies on accurate cellular morphologies, we curated our segmentations to minimize errors. To facilitate curation, we created a custom graphical user interface using napari. In this interface, the analyst can edit individual cells and commit them to the curated segmentation image once they have been validated.

#### Quantifying cellular morphology and tissue organization

Building upon scikit-image and scipy, we developed a pipeline to quantify the morphology of the individual cells. The following metrics were computed for each segmented cell.

##### Volume

We measured the volume of cells by counting the voxels in the cell segmentation.

##### Surface area

To compute the surface area of the cells, we converted the surface of the cell segmentation into a triangular mesh using the marching cubes algorithm. Then we smoothed the mesh using Laplacian smoothing and summed the area of all triangles in the mesh.

##### Elongation

The elongation was calculated using the ratio of the major and minor axis lengths. The major and minor axis lengths were measured using the scikit-image regionprops function. This implementation computes the major and minor axes as the major and minor axes of the ellipse that has the same normalized second central moments as the cell segmentation.

##### Aspect ratio

We computed the aspect ratio of the cells by dividing the cell’s major axis by the mean of the two minor axes. The major and minor axes were measured by fitting an oriented bounding box around the segmentation and computing the length of the edges of the bounding box. The orientation of the bounding box was determined by using principal component analysis to the vertices of the cell meshes to find the principal axes. The major axis was aligned with the principal axis with the highest eigenvalue.

##### Sphericity

The cell sphericity Ψ was calculated based on the measured surface area *A* and volume *V* according to

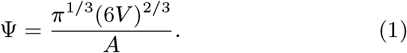

### Inference of mechanical properties with OptiCell3D

#### Mesh generation

OptiCell3D requires multimaterial meshes as input. To generate them from segmentations, OptiCell3D comes with a meshing pipeline that converts instance segmentation label images into triangulated multimaterial meshes. First, an initial mesh is created with the Marching Cubes algorithm, followed by a resampling using Voronoi clustering with PyACVD. For the meshes in this paper, we set the target area of the elements to 2.5 μm^2^. Other geometries may require other target areas. Finally, each element is annotated with the labels of the cells or background that the element is touching.

#### Deformable cell model

For the forward simulation, we represent the tissue patch with a potential energy of the form

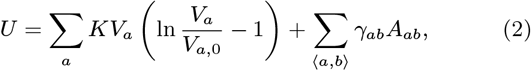

The first term in Eq. 2 accounts for the work done to change the volume of a cell with a given bulk modulus *K* from its equilibrium size *V*_*a*,0_ to *V*_*a*_ [28]. The second term models the contractile tensions generated by the actomyosin cortices. The second sum therefore runs over all interfaces between neighboring cell pairs indexed by *a* and *b*, but also over all (internal or external) tissue surfaces where *b* is then not a neighboring cell, but a free or fixed tissue boundary. *γ*_*ab*_ is the interfacial surface tension. The volume of a triangulated cell reads

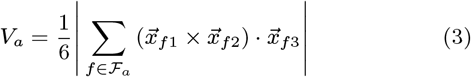

where 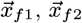 and 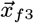 are the nodal coordinates of triangle *f*, and ℱ_*a*_ is the set of triangles of cell *a*. The cellular surface area is calculated as

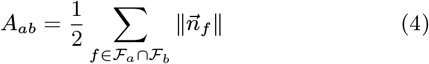

where the sum runs over all triangles making up the interface, and

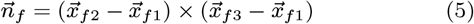

is the unnormalized normal vector of face *f*.

The resulting force on node *i* provides the direction and slope of the steepest descent on the energy landscape:

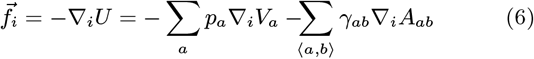

The pressure difference *p*_*a*_ between the cytoplasm of cell *a* and the external medium is given by

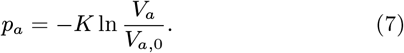

ℱ_*i*_ is the gradient operator with respect to the position of node *i*:

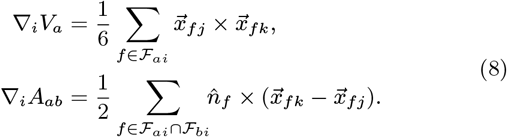

where 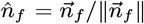 is the unit normal of face *f*, ∇_*ai*_ the set of all triangles containing node *i* on cell *a*, and *i, j, k* are the vertex indices of face *f*.

In the simulations, we solve the overdamped equations of motion for all nodes *i* that are not fixed:

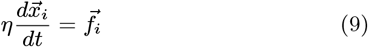

where *η* is a viscosity coefficient that is irrelevant for our method and is set to the arbitrary value of 5 in units of mass per time. The cell nodes are propagated in time with the forward Euler method:

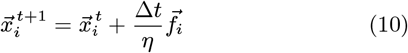

for a total number of *n*_iter_ times.

#### Parameter inference

The loss function that OptiCell3D iteratively minimizes is

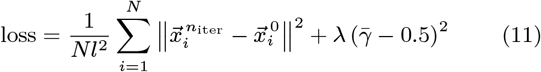

where 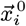 and 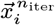 are the initial and final positions of node *i* in the deformable cell model simulation, respectively, *N* is the number of non-fixed nodes, 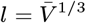 is a characteristic length scale obtained from the average initial cell volume 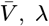, is a regularization coefficient and

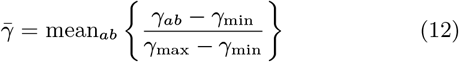

is the mean of all surface tension values normalized to the user-specified range for the initial parameter guess, [*γ*_min_, *γ*_max_]. The second term in Eq. 11 gauges the average surface tension to the midpoint of that range. Note that the simulation is set to a fixed duration and number of time steps *n*_iter_ and does not reach equilibrium. Our method only requires the direction and distance of short-term nodal motion away from the imaged position due to the current mechanical parameter estimate. Minimization is performed with the nonlinear conjugate gradient method. The necessary gradient of the loss function with respect to the parameters is obtained with backward automatic differentiation to match the forward simulation exactly.

#### Implementation

OptiCell3D is provided as a Python library, with the deformable cell model implemented in JAX [29]. Time integration of the forward simulations is performed in Diffrax [39]. The parameters are then updated with the Polak–Ribi`ere conjugate gradient method implemented in the scipy minimize() function [40]. A graphical user interface is provided as a napari plugin.

#### Geometry perturbation pipeline

We developed a geometry perturbation pipeline to simulate the inaccuracies in cell boundary positioning commonly introduced by segmentation tools, and to evaluate their impact on parameter inference. It starts by generating a 3D Perlin noise image [30] with 125 voxels along all axes (Supplementary Fig. S3a). The noise frequency can be adjusted by controlling the number of octaves used in generating the Perlin noise. Noise values at each voxel are normalized to lie within the interval [− *d*_max_, *d*_max_], where *d*_max_ determines the intensity of the perturbation. Next, the mesh of a simulated geometry is overlaid and its nodes are displaced in normal direction based on the local noise intensity (Supplementary Fig. S3b). The resulting perturbed mesh is then converted into a segmentation label image with a voxel size of (0.25 μm)^3^. Examples of perturbations with different intensities *d*_max_ and number of Perlin octaves are shown in Supplementary Fig. S3c.

#### Hyperparameter tuning

OptiCell3D has five hyperparameters (Table 2). To optimize them, we first generated *n* = 10 geometries in SimuCell3D [28], each containing 16 cells resembling early embryos. These geometries were then perturbed using the above-mentioned pipeline with varying Perlin noise patterns. We then employed Latin hypercube sampling to draw 10,000 hyperparameter sets with bounds listed in Table 2. OptiCell3D was run with each set of sampled hyperparameters on the perturbed geometries. The 200 hyperparameter sets with the lowest mean sum of pressure and surface tension estimation errors are shown in Supplementary Fig. S7a. Next, we varied each hyperparameter from this initial selection individually, keeping the others fixed at values near their optimum (Supplementary Fig. S7b–g). This approach allowed us to estimate the optimal values for each hyperparameter as listed in Table 2.

#### Accuracy Benchmarks

For the inference accuracy benchmarks, we chose to start with segmentations and not microscopy images, as the primary goal of each of the tested methods is parameter inference and not segmentation. Further, different types of images and samples will likely require different segmentation methods. For each inference method we used the meshing pipeline recommended by the method’s documentation. We created equilibrium tissue geometries in SimuCell3D [28] with known cell pressures and interface surface tensions drawn from a uniform distribution over (2.8, 5.8) ×10^−4^ N/m. To benchmark Foambryo, we installed version 0.3.5 and performed meshing and inference using default parameters. For the FIDES benchmarks, we downloaded FIDES from the Bitbucket repository [19] (SDT-PICS commit hash: 3760ab9, FIDES commit hash: fd339ae). For “Liu 2023”, we obtained the implementation from the official GitHub repository at commit 5dd87e9. For each estimated pressure and surface tension parameter, we computed its relative error as 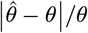, where 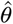 represents the estimated value and *θ* the true value.

#### Power analysis

We estimated the number of samples required to resolve differences between parameters ranging from 1% to 20% by simulating the comparison of two sets of unpaired samples. To replicate the theoretical measurement distributions of two parameters, we first drew two sets of inference errors from the noise-free error distributions of each pipeline (Supplementary Fig. S1). Then, we shifted these error sets based on the tested effect size and used the Mann–Whitney U test to compute the probability that the two simulated measurement distributions were statistically different. This analysis was repeated 10,000 times, each with different random draws from the inference error distributions.

#### Runtime benchmark

We measured the wallclock time required by each pipeline to first convert an input image with a voxel size of (0.25 μm)^3^ into a mesh, and to then estimate the cell parameters from this mesh. Segmentation label images were used as input for both Foambryo, “Liu 2023” and OptiCell3D. For FIDES, we first converted grayscale images into meshes using the SDT-PICS meshing tool [41]. In these images, voxels had a value of 255 at cell boundaries and 0 elsewhere. All computations were performed on an Intel Xeon W-2125 processor (4 cores, 4.0 GHz).

#### Statistical analysis of distribution differences

The surface tension and pressure distributions inferred for a given tissue by OptiCell3D are normalized to a constant mean, rendering standard statistical tests inappropriate and prone to spurious p-values. To account for the compositional nature of these data, we first applied a centered log-ratio (CLR) transformation to all samples. Statistical differences between distributions, both within and across samples, were then assessed using permutation tests. Significance was evaluated using a two-sided permutation framework with 10,000 independent resamples.

#### Forward simulations of epithelial tissues

All simulations of epithelial tissues were performed using the SimuCell3D framework. Details of its implementation are described in [28]. The simulations were initialized with a three-dimensional honeycomb lattice containing 93 cells. At the start of each simulation, all cells had identical heights, and the apical, lateral, and basal surface tensions were uniform across all cells within a given simulation. Initially, all cells were in contact on their basal side with a static support, modeled as a special immobile cell. Cells were allowed to slide freely along this support throughout the simulation. The cells did not grow nor divide during these simulations but simply deformed until reaching an equilibrium shape. Across different simulation runs, the apical surface tension was varied, while all other parameters were kept constant. Each simulation was run until the system reached mechanical equilibrium. Cell heights were then quantified by measuring the length of the apico-basal axis of each cell.

## Code Availability

OptiCell3D is open source and publicly available from the date of publication at https://git.bsse.ethz.ch/iber/Publications/2026_runser_opticell3d. The software is distributed under the BSD 3-Clause License.

## Data Availability

The raw data generated and analyzed in this study will be made publicly available from the date of publication.

## Acknowledgements

We thank Dimitri Spicher and Hannes Welle for preliminary work and valuable discussions. This work was funded by Swiss National Science Foundation project grant no. 219990 to DI.

## Competing Interests

SR, FC, LC, LS, RV, and DI declare no competing interests. KY, MA, and FLL are currently employed in private sector positions. Their contribution to this study occurred exclusively while employed at ETH Zurich. The opinions expressed in this manuscript are those of the authors and do not necessarily reflect the views of the employer.

## Author Contributions

Concept OptiCell3D: SR, MA. Model and algorithm development: SR, KAY. Hyperparameters screen: SR, MA. OptiCell3D accuracy benchmarking: SR. Image acquisition: LS, LC, FLL. Segmentation: KAY, FLL, FC. Morphometrics: KAY, SR, FLL, FC. Biological Parameter Inference: KAY, SR. Figures: SR, KAY. Writing: SR, KAY, RV, DI. Supervision: RV, DI. Funding acquisition & overall concept: DI.

