## Supplementary Tables and Figures for "OptiCell3D: Precise inference of mechanical cell properties from microscopy imaging"

### Supplementary Information

**Table 1:** Feature comparison of image-based force inference methods.

| Framework name | Publication year | Dimension | Viewer | Code available | License | Reference |
|---|---|---|---|---|---|---|
| VFM | 2010 | 2D | No | No | — | [8] |
| MI | 2012 | 2D | No | No | — | [9] |
| — | 2012 | 2D | No | No | — | [10] |
| CellFit-2D | 2014 | 2D | Yes | Yes | — | [11] |
| CellFit-3D | 2017 | 2.5D | No | No | — | [12] |
| — | 2018 | 3D | No | No | — | [17] |
| VMSI | 2020 | 2D | Yes | Yes | — | [13] |
| — | 2022 | 2D | No | Yes | MIT | [16] |
| Foambryo | 2023 | 3D | Yes | Yes | CC BY-NC-SA 4.0 | [18] |
| FIDES | 2023 | 3D | No | Partial | — | [19] |
| — | 2023 | 3D | No | Yes | — | [20] |
| ForSys | 2023 | 2D | Yes | Yes | BSD-3 | [14] |
| — | 2024 | 3D | No | Yes | CC BY-NC-SA 4.0 | [21] |
| ForceIn3D | 2025 | 3D | No | No | — | [23] |
| Vertax | 2026 | 2D | No | Yes | CC BY-SA 4.0 | [15] |
| OptiCell3D | 2026 | 3D | Yes | Yes | BSD-3 | this work |

**Table 2:** Hyperparameters used for the optimization in OptiCell3D.

| Parameter | Description | Tested range | Selected value |
|---|---|---|---|
| $n_{\text{iter}}$ | Number of iterations to perform in the forward simulation in each optimization loop | 1 – 250 | 25 |
| $\Delta t$ | Size of the time step in the forward simulations, in seconds | 1 – 1000 | 170 |
| $\lambda$ | Regularization coefficient in the loss function | $10^{-6}$ – $10^{12}$ | $10^{3}$ |
| $n_{\text{start}}$ | Number of optimization runs from which the parameters with the lowest loss are selected | 1 – 100 | 5 |
| $n_{\text{opt}}$ | Number of optimizer iterations performed per run | 1 – 1000 | 50 |

**Table 3:** Quantitative morphological characteristics of different tissue types. Values are reported as mean ± SD.

| Tissue type | Number of cells | Cell area (μm$^2$) | Cell volume (μm$^3$) | Sphericity | Aspect ratio |
|---|---|---|---|---|---|
| Bronchiole | 139 | $2.8 \times 10^3 \pm 9.1 \times 10^2$ | $1.0 \times 10^4 \pm 4.4 \times 10^3$ | $0.80 \pm 0.05$ | $2.0 \pm 0.4$ |
| Kidney | 21 | $4.0 \times 10^2 \pm 5.6 \times 10^1$ | $5.6 \times 10^2 \pm 1.5 \times 10^2$ | $0.81 \pm 0.03$ | $2.1 \pm 0.3$ |
| Intestine | 237 | $1.9 \times 10^3 \pm 4.2 \times 10^2$ | $4.7 \times 10^3 \pm 1.3 \times 10^3$ | $0.72 \pm 0.05$ | $3.6 \pm 0.8$ |
| Esophagus | 75 | $2.9 \times 10^3 \pm 1.1 \times 10^3$ | $1.1 \times 10^4 \pm 6.0 \times 10^3$ | $0.81 \pm 0.03$ | $2.2 \pm 0.4$ |
| Bladder | 739 | $9.9 \times 10^2 \pm 4.4 \times 10^2$ | $2.2 \times 10^3 \pm 1.5 \times 10^3$ | $0.80 \pm 0.05$ | $2.7 \pm 0.8$ |

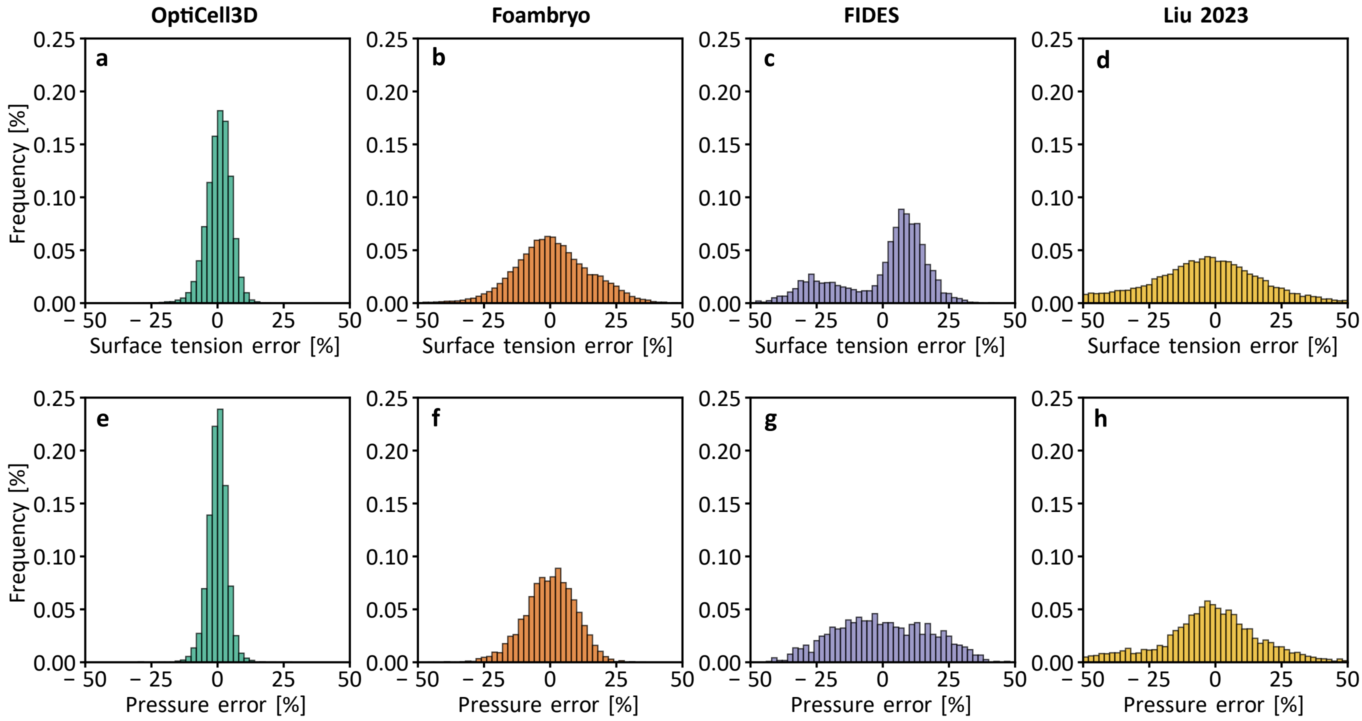


**Figure S1: Empirical distributions of parameter estimation errors. a–c,** Distributions of surface tension estimation errors from OptiCell3D, Foambryo, FIDES, and “Liu 2023” on $n = 250$ simulated geometries, with 62,450 individual interfaces estimated for Foambryo, “Liu 2023”, and OptiCell3D, and 8844 for FIDES. **d–f,** Distributions of pressure estimation errors on $n = 250$ simulated geometries, with 6200 cell pressures estimated for Foambryo, “Liu 2023”, and OptiCell3D, and 1176 for FIDES.

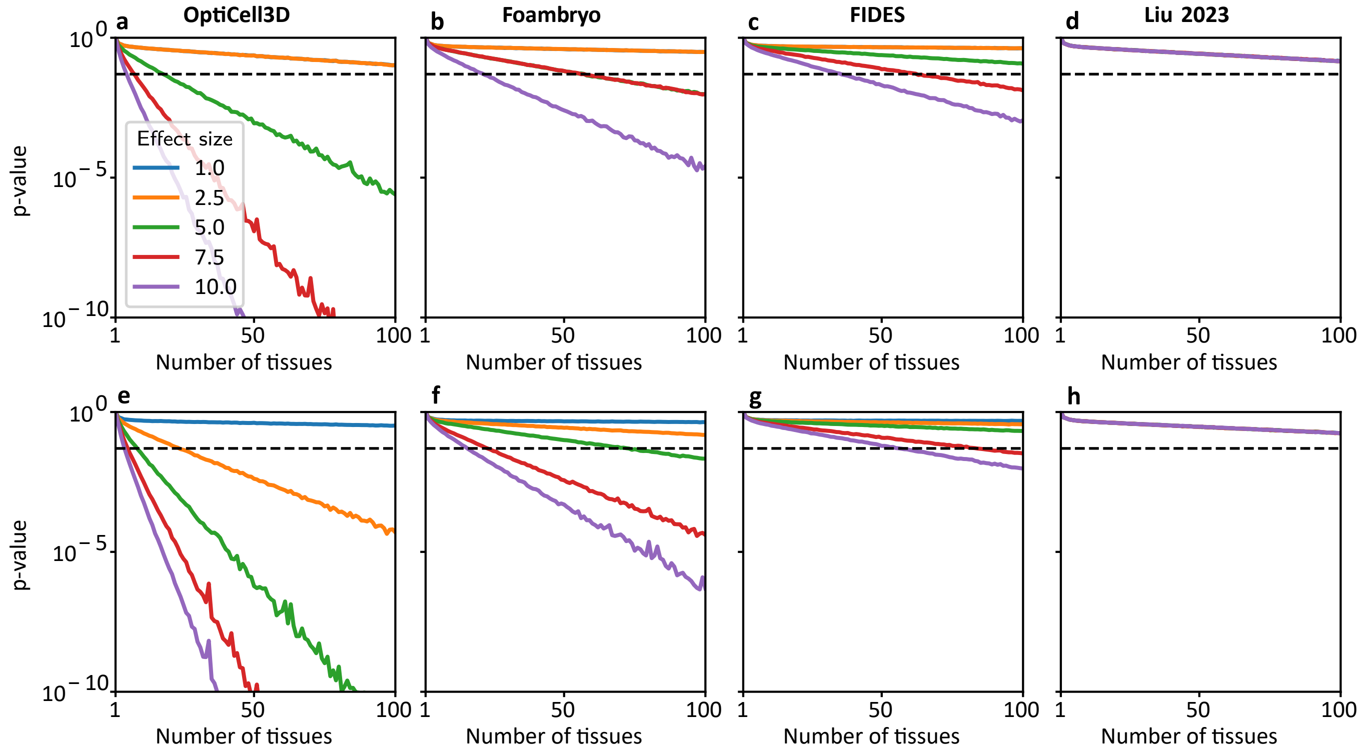


**Figure S2: $p$-values of differences between measurement distributions. a–c,** Mann–Whitney U $p$-values of two distributions of surface tension measurements shifted by a given relative difference (colors) for a given number of tissue samples. **d–f,** Analogous $p$-values of pressure measurements. Black dashed lines mark $p = 0.05$. All plots use $n = 10{,}000$.

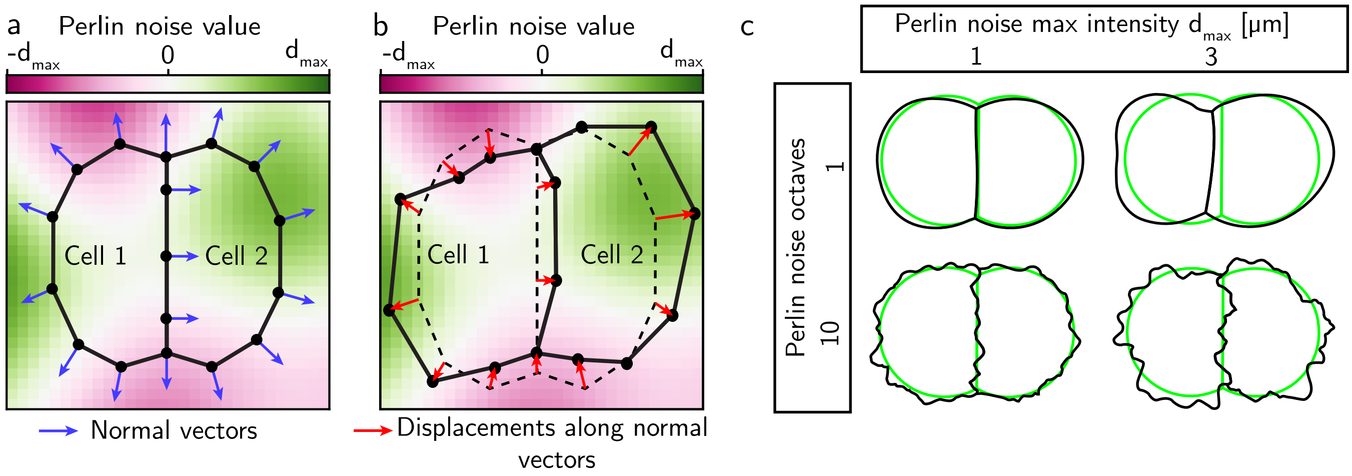


**Figure S3: Geometry perturbation procedure.** For clarity, the three-dimensional perturbation process is illustrated in two dimensions. **a,** The spatial domain is discretized into a uniform grid of Perlin noise values (colors) in the interval $[-d_{\max}, d_{\max}]$. **b,** Mesh nodes are displaced along their surface normals in proportion to the local noise values. **c,** Examples of perturbed geometries (black) compared to their original shapes (green) as a function of the noise profile.

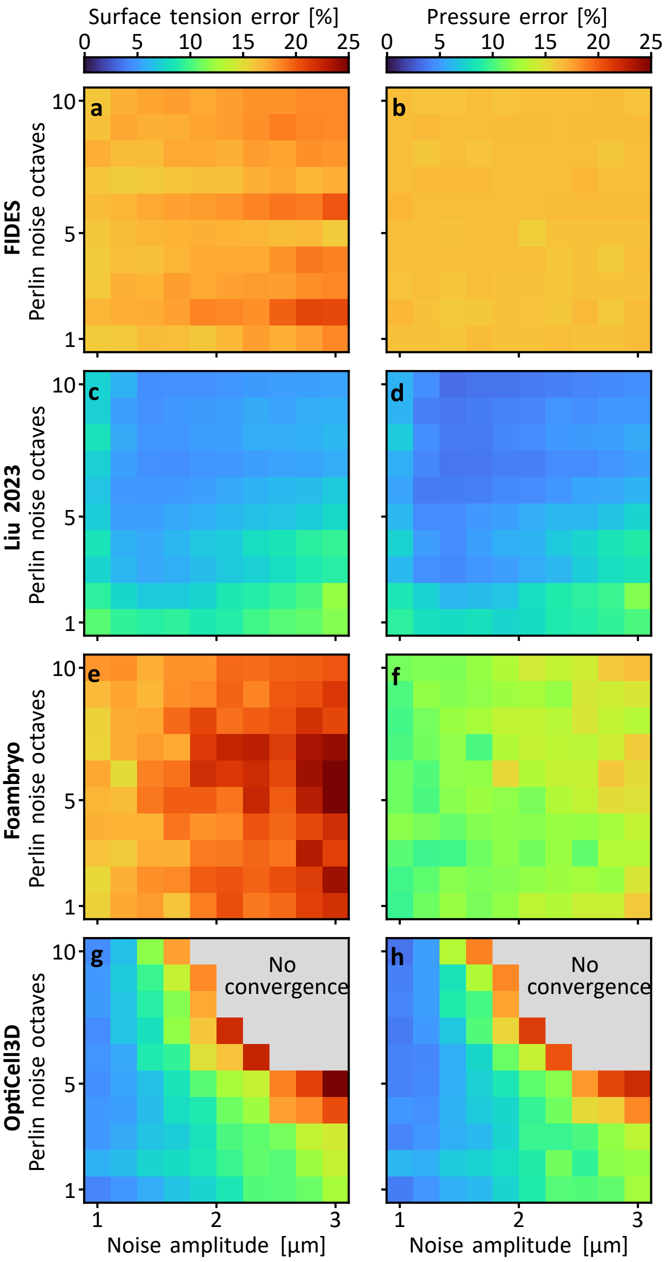


**Figure S4: Parameter estimation error due to geometric perturbations. a–h** Mean relative surface tension and pressure error for FIDES (**a,b**), "Liu 2023" (**c,d**), Foambryo (**e,f**), and OptiCell3D (**g,h**). Grey regions in panels g and h indicate noise profiles where loss values exceeded a maximum threshold of $2.5 \times 10^{-4}$. All plots use $n = 10$ 8-cell geometries.

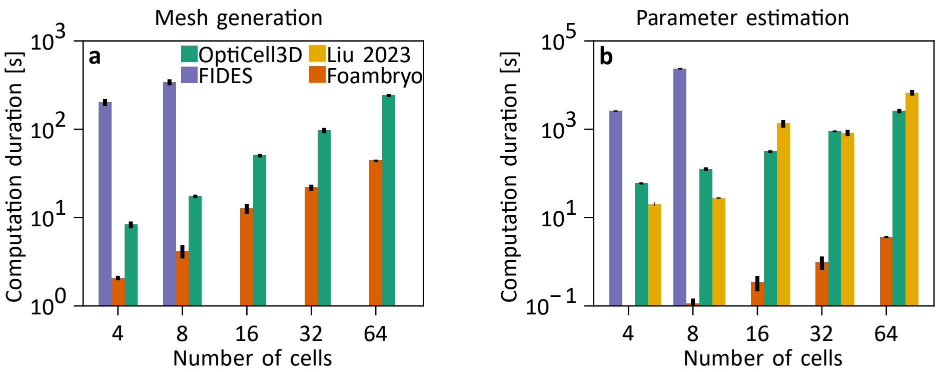


**Figure S5: Computational performance benchmark.** **a,** Computation time required to convert an image of a tissue with voxel size $(0.25\,\mu m)^3$ into a mesh as a function of the number of cells. **b,** Computation time required to estimate the cell parameters from an input mesh as a function of the number of cells. The duration measurements of FIDES with 16, 32, and 64 cells were omitted due to prohibitively long runtime. All computations were performed on an Intel Xeon W-2125 processor (8 cores, 4 GHz). Bars are means ± SD over $n = 5$ tissues.

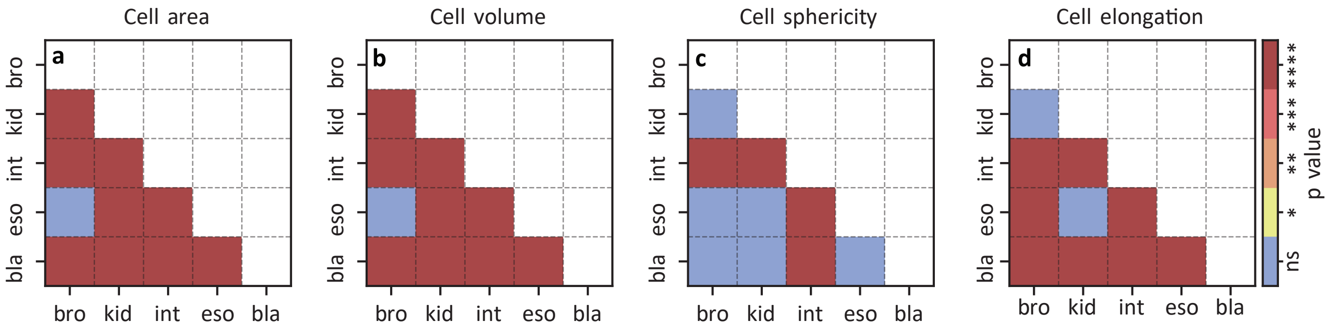


**Figure S6: Pairwise comparison of morphometric feature distributions.** For each analyzed morphometric feature, pairwise differences in distribution means between all tissue combinations were evaluated using the Mann–Whitney U test. $p$-values were adjusted for multiple comparisons using the Benjamini–Hochberg false discovery rate procedure.

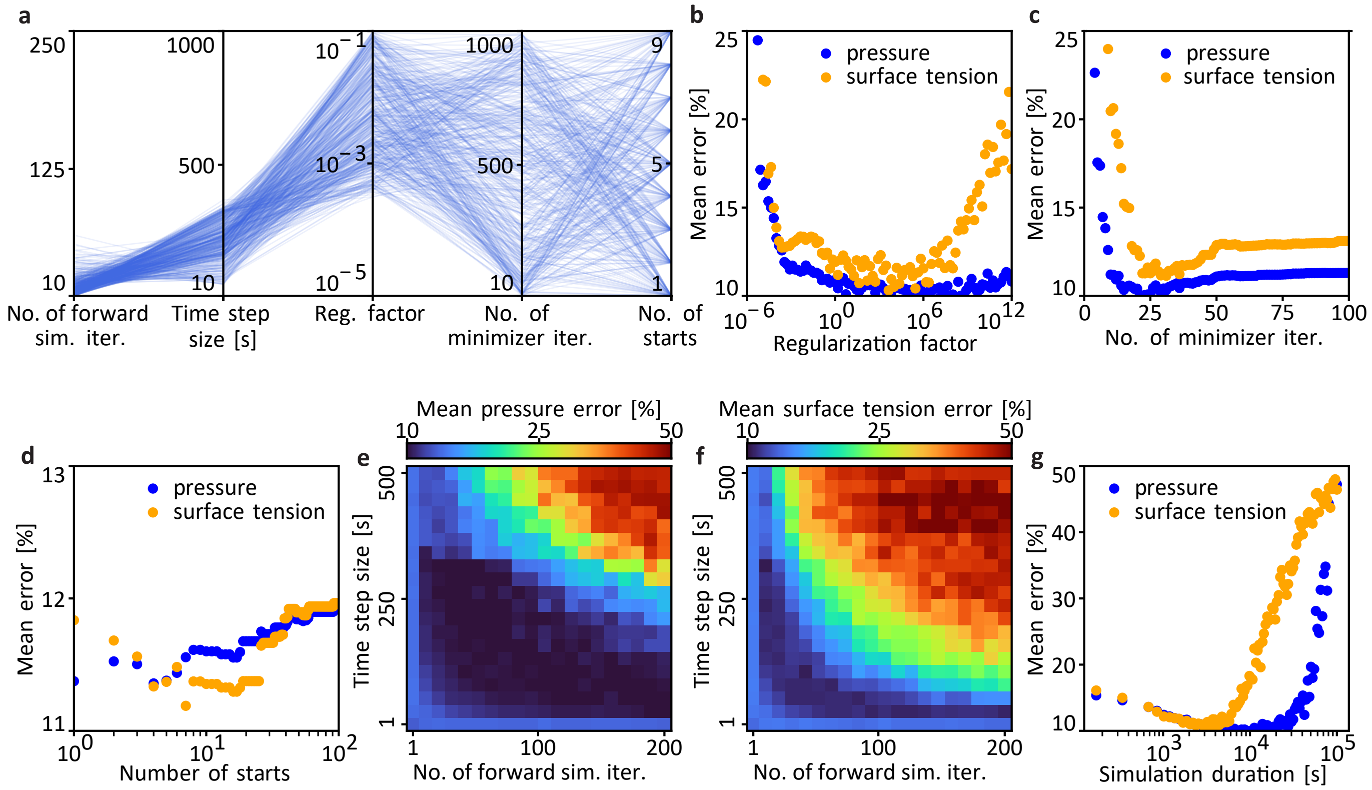


**Figure S7: Hyperparameter optimization.** **a,** Parallel coordinate plot showing the 200 out of 10,000 hyperparameter sets (lines) with the lowest combined mean surface tension and pressure errors. **b,** Mean relative estimation error as a function of the loss regularization factor, $\lambda$. **c,** Mean relative estimation error as a function of the number of minimizer iterations, $n_{\text{opt}}$. **d,** Mean relative estimation error as a function of the number of runs in the multi-shoot approach, $n_{\text{start}}$. **e,f** Mean relative parameter errors as a function of the number of forward simulation iterations, $n_{\text{iter}}$, and time step size, $\Delta t$. **g,** Mean relative estimation as a function of the forward simulation duration, $n_{\text{iter}}\Delta t$. All plots use $n = 10$ geometries.
